# Non-destructive spatial mapping of vegetation plots: a re-introduction of the pantograph

**DOI:** 10.1101/2023.04.28.538692

**Authors:** Trace E. Martyn, Margaret M. Mayfield

## Abstract

1. Non-destructive spatial mapping of herbaceous plants is often not possible with many modern imaging techniques, especially in systems with highly structured, dense herbaceous canopies. For this purpose we suggest using a modern version of the classic pantograph, a simple instrument that allows precisely scaled drawings. The pantograph version we describe here was specifically designed for small-scale herbaceous vegetation mapping.
2. Specifically, our pantograph design is useful for rapidly collecting accurate, spatially explicit data at the scale of 0.1-2 m^2^ and includes a paired drawing board for easy use in field conditions.
3. We tested the design and technique on 100 annual plant plots that ranged in total density and in plant stature. Based on this mapping trial, we present guidelines for effective manual mapping and map digitization.
4. A pantograph is a useful and inexpensive to make tool for non-destructively spatially mapping individual herbaceous plants in the field. Here, we present instructions for the design and fabrication of our modern pantograph, board, and pencil attachment designed specifically for researchers wanting to include small-scale spatial context in their research.

## Introduction

There is widespread interest in understanding the role of spatial context in community ecology. At broad scales, remote sensing has become the go-to technology to identify plant species and rapidly and accurately estimate percent cover (Gould, 2000; Kwok, 2018). Satellite/image-based technologies often, however, rely on ground surveys to accurately pinpoint exact plant locations given the issues associated with dense canopies (i.e. stem/trunk locations). Within forests, stem locations are often easy to assess using a GPS or basic X-Y coordinate planes (or more recently LiDAR; Balsi *et al*. 2018) but for herbaceous vegetation it can be harder to delineate individual plants with these methods. Here, we re-introduce the pantograph for this purpose. The pantograph is a classic ecological survey device and is scaled well for surveys of herbaceous plant communities. In this paper we present plans for the construction of such device using modern materials and tools. We also offer insights and details about how this simple tool can aid in a wide range of ecological projects on herbaceous plant communities.

The pantograph was originally designed by Christoph Scheiner in 1631 (Scheiner, 1631) and is used to copy, reduce or enlarge subjects based on scaling ratios and similar triangles. This technique was popular in the early- to mid-20th century in the Western United States ecology community (Bakker *et al*., 2008; Hill, 1920) and has been primarily used to map vegetation in desert, grassland, and shrubland ecosystems including as part of the Long-Term Ecological Research (LTER) program (see data papers Adler *et al*. (2007); Anderson *et al*. (2012, 2011); Chu *et al*. (2013); Zachmann *et al*. (2010)). In these studies, the pantograph was used to understand long-term changes in vegetation (Albertson & Tomanek, 1965), patterns of survivorship (Chu *et al*., 2014), demography (Chu & Adler, 2014; Dalgleish *et al*., 2010; Gong *et al*., 2011; West, 1979), and coexistence/diversity (Adler *et al*., 2003; Chu *et al*., 2013). Though still an effective tool, classic pantograph techniques can be time consuming and two people are generally required to implement them (Ellison, 1942; Hill, 1920; Pearse *et al*., 1935). In this paper, we present an updated design and technique that is easily performed quickly by a single person. Our technique is particularly effective for questions requiring data on specific spatial orientation of individuals in high density plots where plant stems are close together (Ellison, 1942; Hanson & Love, 1930).

Other modified pantograph methods have been developed in recent years that streamline the pantograph and combine mapping and digitalization processes. Wallin and Avery (Wallin & Avery, 2007) for instance, designed a frame with an electronic probe that would identify an individual’s location on an existing map of the plot based on stem location relative to sensors placed in the corner of plots. This method is quick and easy and is especially useful for repeatedly locating specific individual plants within field plots (Wallin & Avery, 2007). It has not, however been tested as an approach for mapping entire communities and A disadvantage of this approach is the cost associated it. The digital equipment may be expensive for students or researchers working on a tight budget. Other recent approaches to small scale mapping for estimating percent cover and biomass and identifying specific species or individuals have been developed using digital photography and/or spectral imaging (Klauschies *et al*., 2016; Schmidt & Skidmore, 2003; Wilson, 2011). These techniques are also costly and limited to locations for which images are available (Mumby *et al*., 1999). These techniques are also ineffective for high density plant communities where overlapping plants make defining the actual rooting location indiscernible from an image.

### Aims

We present a detailed design and trial of our field pantograph including details on various scaling ratios, assembly techniques, materials, and considerations for using a pantograph for ecological mapping.

## Methods

Our pantograph design is made up of two components: an “arm” and a drawing board (Figure 1). We present our design and suggestions for both the arm and board as well as an approach for digitizing the maps and for analysis of spatial data collected using the pantograph.

**Figure 1:**
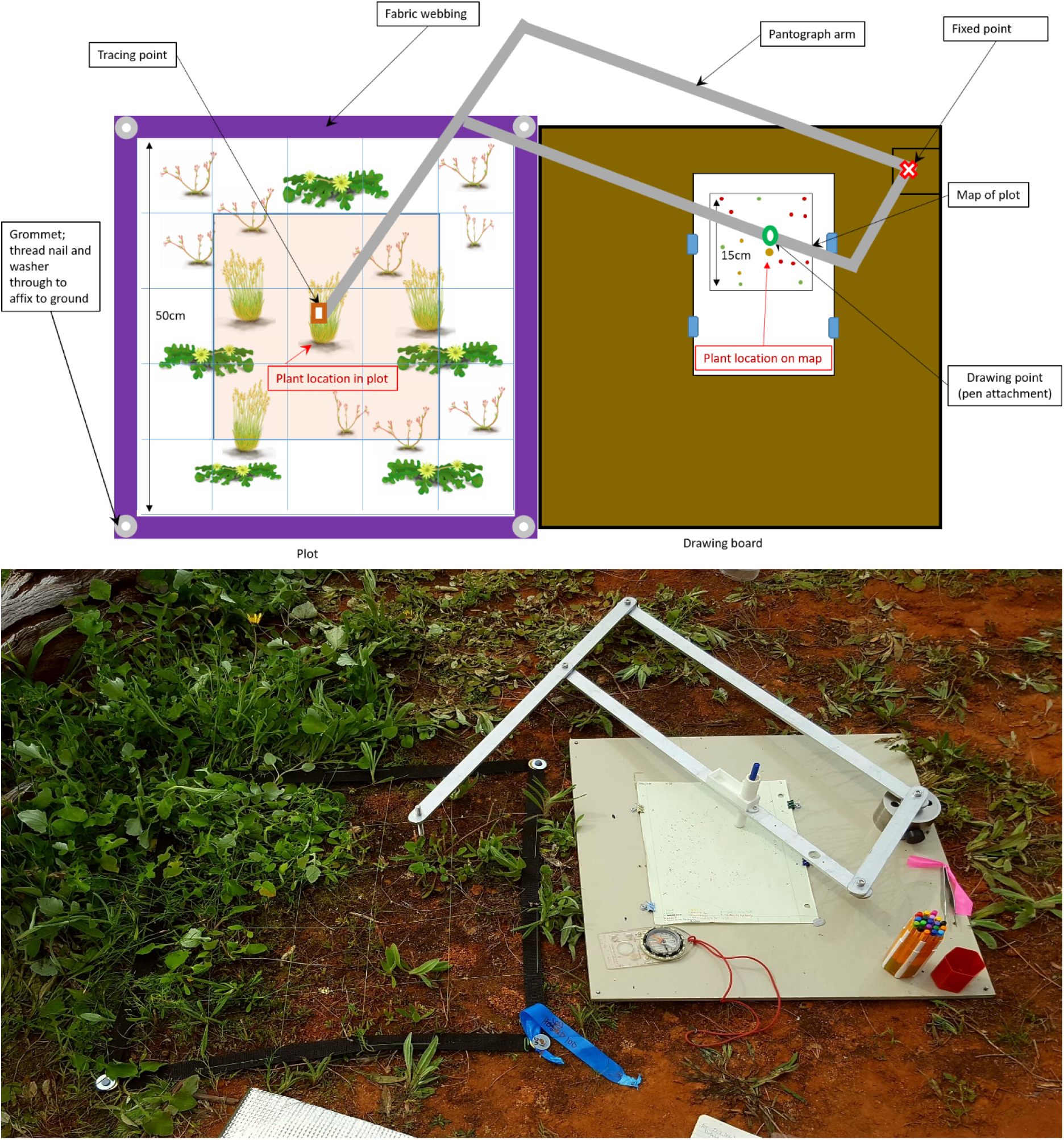
Pantograph layout design and in action. The top image shows a diagram of the pantograph in use next to a plot. The blue lines in the plot represent a 10 cm grid that was placed over the plot and the orange shaded section indicates the area of focus with the surrounding white section is the spatial buffer zone. The bottom image shows the pantograph in use in the field in York gum-jam woodlands in Western Australia.

### Pantograph arm design

The pantograph arm follows a classic design based on scaling ratios and similar triangles (Scheiner, 1631). As the design described here is easily scalable to specific projects, the first step is to identify the reduction ratio that works for the system and application of interest. The reduction ration is how much you need the pantograph to scale down your plot so that your detailed map will fit onto a sheet of paper (Figure 1, Table 1). The scaling ratio determines the length of the longest arm of the pantograph (A in Figure 1). We matched the length of the longest arm with the length of one edge of our plot plus and extra 10 cm to provide flexibility in board placement on the ground (instead of being restricted to placing the board exactly next to the plot). The arms can be assembled in two ways depending on where the user wants the drawing point to be; we use the arrangement shown in Figure 2b)

**Table 1:** Example lengths for pantograph arm construction for three different vegetation plots sizes.

| Ratio | Section Parts (cm) |  |
| --- | --- | --- |
|  | A | B |
| 30 X 30 cm plots |  |  |
| Longest arm = 40 cm |  |  |
| 2 : 1 | 20.0 | 20.0 |
| 8 : 3 | 25.0 | 15.0 |
| 3 : 1 | 26.7 | 13.3 |
| 4 : 1 | 30.0 | 10.0 |
| 50 X 50 cm plots |  |  |
| Longest arm = 60 cm |  |  |
| 3 : 1 | 40.0 | 20.0 |
| 4 : 1 | 45.0 | 15.0 |
| 5 : 1 | 48.0 | 12.0 |
| 6 : 1 | 50.0 | 10.0 |
| 100 X 100 cm plots |  |  |
| Longest arm = 110 cm |  |  |
| 5 : 1 | 80.0 | 20.0 |
| 6 : 1 | 83.3 | 16.7 |
| 10 : 1 | 90.0 | 10.0 |

Note: Need to put some info here...

**Figure 2:**
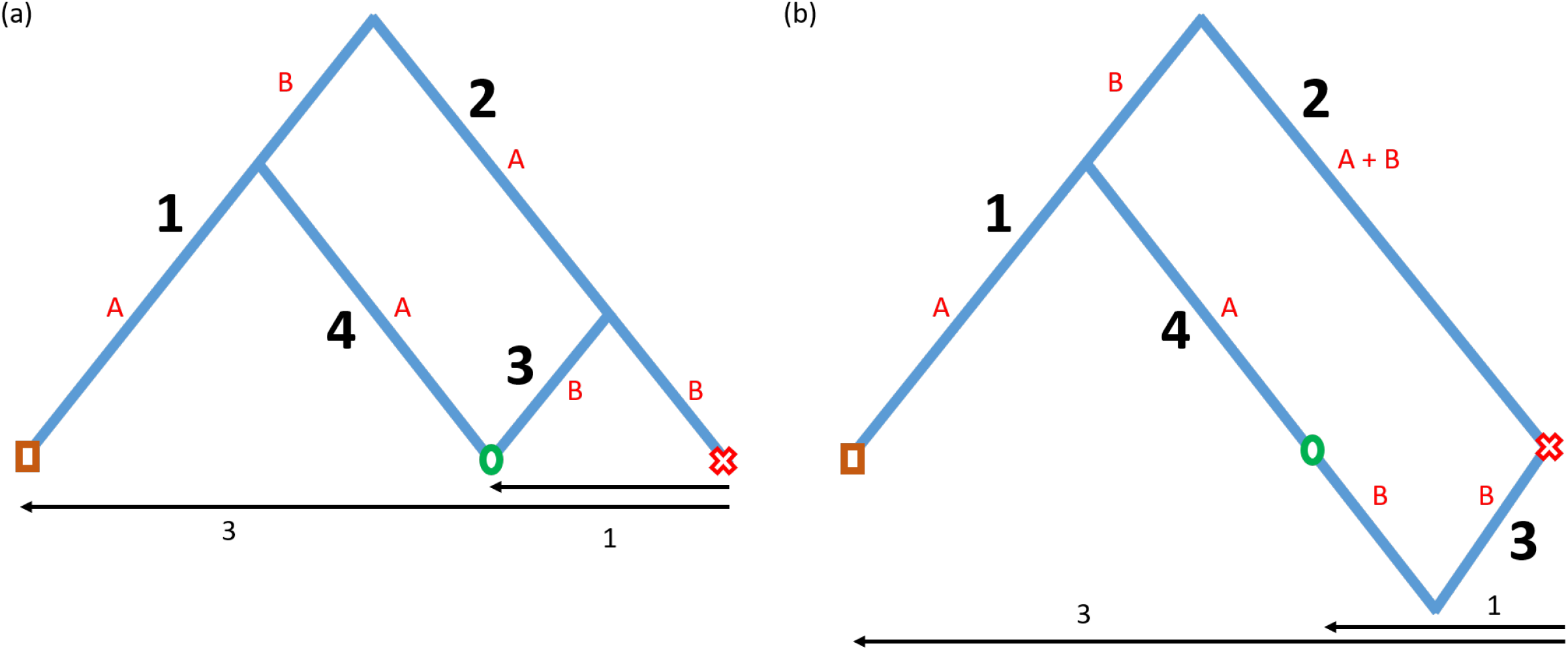
Two different ways to arrange pantograph. Different ways to assemble the pantograph. Both methods will create identical maps, the difference is the placement of the drawing point (green circle), either at the hinge joint (a) or within the length of Section 4 (b). Pantograph arm sections are labelled 1-4 with bold text and diagrammed further in Figure 3. Red letters (A and B) relate to relevant parts of each arm section where the ratio of these parts match the intended reduction ratios (see Table 1). Tracing point (brown square) and fixed point (red “X”) are also represented on the figure. We diagram the pantograph arm, here, to show how the rules for similar triangles can be used with the intended plot to map reduction ratios (here representing and 3:1 reduction through the arrows at the bottom of the plot).

For the arm, we used a 5 mm thick x 25 mm wide piece of aluminum. This allowed the arm to be durable, usable in wet weather and light enough to carry into the field by a single person. Each section of the arm is attached with a hinge joint designed to hold the arms together while allowing them to move freely. Our design uses a nylock nut, a nylon washer, and a hex screw at each of the four hinge points (purple, orange, green, and red points in Figure 3; assembly diagram, Figure 4). The pantograph arm is made of four cut pieces (Figure 3) of this aluminum with lengths and locations of hinge points decided using an equation for similar triangles. Below are the calculations for arm sections A and B (see Figure 2 and 3) for pantograph arm with longest section length of 60 cm to reduce a 50 × 50cm plot with a 3:1 reduction ratio:

**Figure 3:**
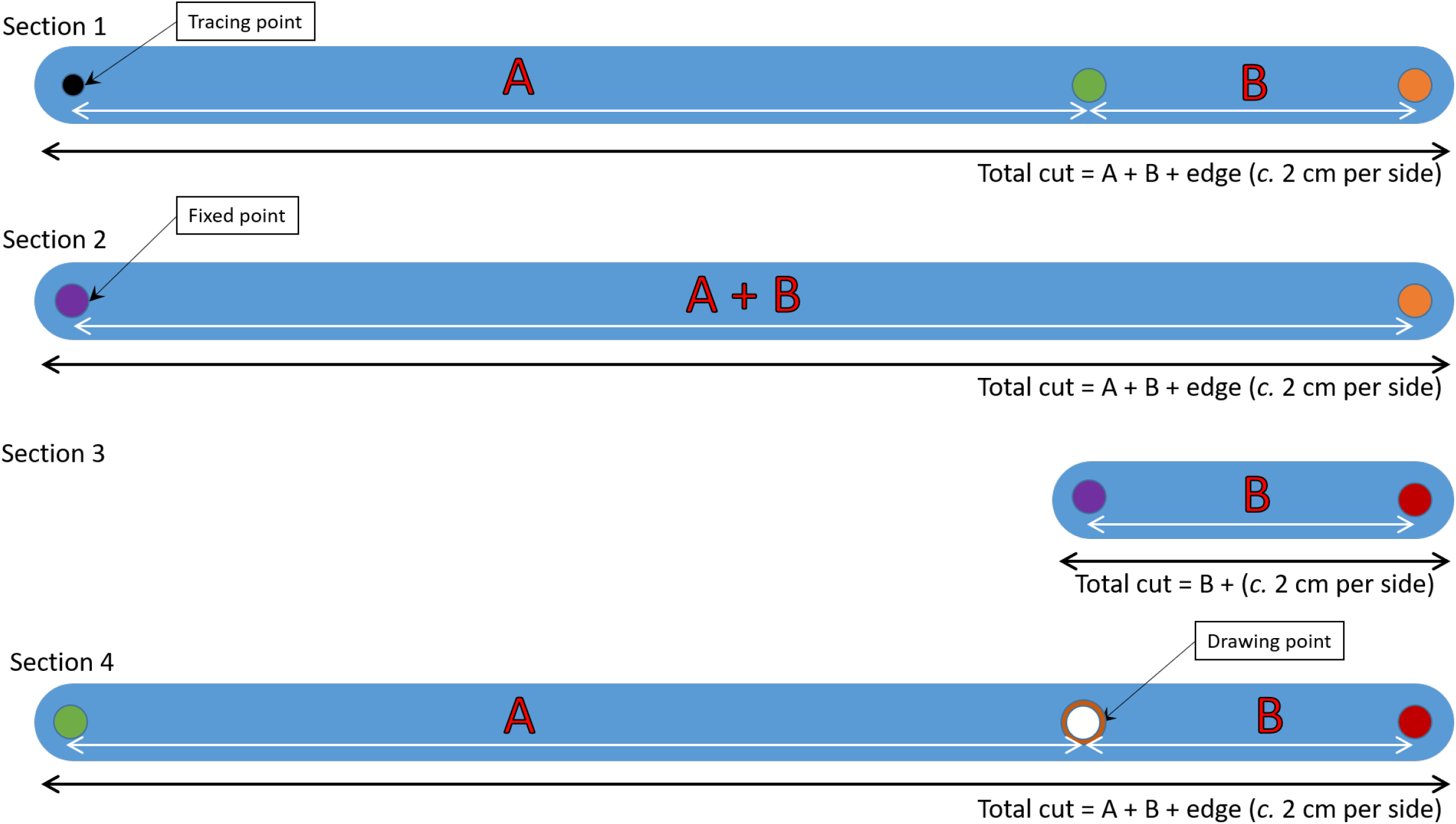
General pantograph arm lengths. Lengths of each segment of the pantograph referencing assembly method in Figure 2. Values for the segments A and B are the distances between each hinge or specific point and examples can be found in Table 1 for different reduction ratios. Red, orange, and green points in the above figure would be matched and threaded together using a screw, nylon washer, and a nylock nut (see Figure 4). These represent the movable points on the pantograph. Total segment value would be slightly greater than the actual ratio calculations so that there is length at the ends of the segments for assembly (see calculations below each section adding *c*. 2 cm per side (4 cm total) to the total cut length to allow for smoothing of edges).

**Figure 4:**
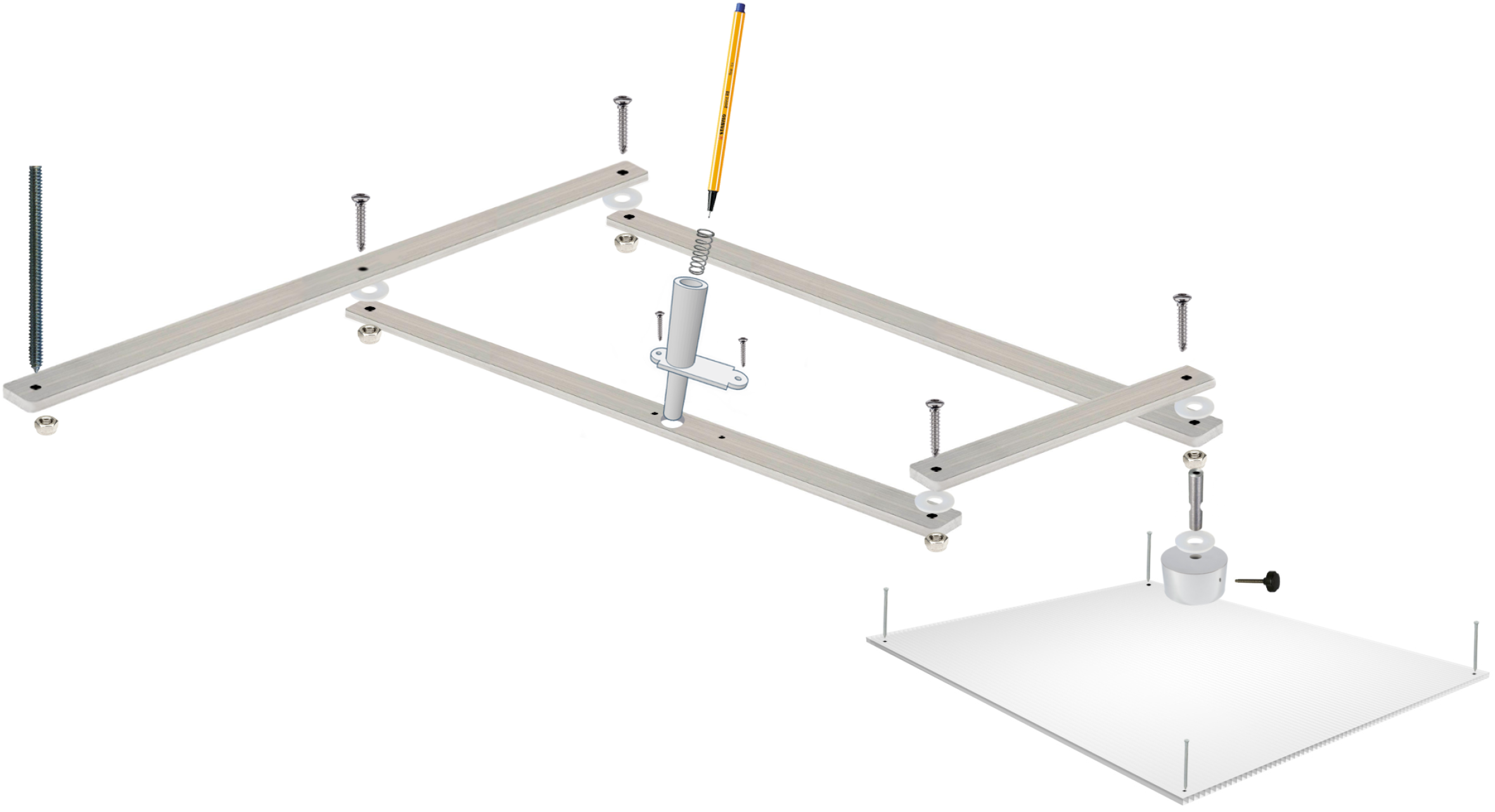
General pantograph assembly. Diagram of the pantograph arm and board assembly. The four segments of the arm are assembled using a screw, nylon washer, and nylon nut. The arm is mounted on the board at the fixed point (metal cylinder) that is affixed to the drawing board by three metals screws through the bottom of the board into the metal piece (not shown). The board is made of a sturdy material (here PVC) and nailed until the ground at the four corners. Images in diagram are not to scale and are not exact representations of materials used.

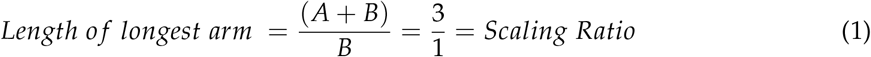

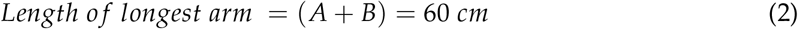

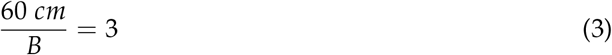

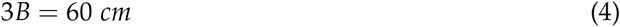

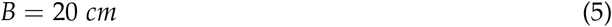

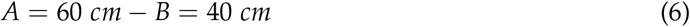

In Table 1, we present four different scaling ratios for common herbaceous plant quadrat sizes (30 cm, 50 cm, and 100 cm squares) to a standard A4 or US Letter piece of paper.

We attached the pantograph arm to the drawing board at a fixed swivel point that includes a weighted block of metal (to counter the weight of the arm and reduce strain on the board). We placed the fixed point towards the top of the drawing board on the board edge opposite the plot (Figure 1 and 4. At the fixed point (purple Figure 3) arm, we added a notched metal piece that threads onto the screw to make the hinge joint (Figure 4). This metal piece slid into a hole of slightly larger diameter drilled into the weighted block on the board. Perpendicular to the larger hole was a smaller threaded hole which was drilled into the side of the weighted block. This smaller hole allowed a wing screw or knob screw to be inserted and fit within the notch in the metal piece (Figure 4). This prevented the arm from popping free of the fixed metal points but allowed the arm to swing freely.

The tracing point on the arm can be any adjustable fine tipped stylus. We recommend a sharpened threaded rod that can be easily adjusted with a nut as needed (Figure 4).

We designed a pen attachment for our pantograph so that the device can be used easily by one person. We used a basic free online 3D graphics program (Tinkercad; Autodesk 123 (2018)) to create a pen attachment that would hold our pens and printed it in plastic using a standard 3D printer (Appendix A, Figure A1). To allow the stylus to move up and down, we used a spring which fit within the inner diameter of the attachment and cut it to the right length so that the tip of the pen did not extend past the tube but when depressed the pen emerged from the tube and marked the paper. We then added the pen attachement to the arm at the drawing point of the pantograph using two small screws (Figure 4; Appendix A, Figure A1).

### Drawing board design

The drawing board can be made of any material and should be selected for the particular environment and project. For our test study, we used a 50 × 50 cm square of PVC so that the board would be waterproof and would lie flat on the ground. To secure the board to the ground, we drilled holes into the four corners of the board (Figure 4). In the field, we then hammered nails through the board into the ground to assure the board remains in the same location throughout the mapping process. We attached the paper to the board using small binder clips screwed into the board and poster tack to prevent the paper from swivelling (Figure 1.

### Mapping protocol

We used our pantograph to study the spatial context on plant-plant interactions in an annual wildflower community in Southwest Western Australia. In this paper, we present details associated with the mapping of these communities to illustrate the types of outcomes possible using this pantograph design. Before mapping the vegetation in our study, we installed fabric plot frames with a 10 cm fishing wire grid around the vegetation. This frame split each 50 cm x 50 cm plot into 25 subsections which helped keep track of which sections we had mapped (Figure 1). We placed our frames approximately six weeks prior to peak biomass to allow the vegetation to grow up around the frames but the pantograph approach does not require that the frame to be set in advance, it would work fine in a plot marked established at the time of sampling. Before mapping we placed the drawing board beside each plot and attached the pantograph arm to the board, nailing the corners down once positioned well. Some fiddling is required to ensure the board is close enough to the plot for the long arm to reach the farthest corner and make a mark on the paper. To insure a stable drawing surface on rocky or uneven ground, adjustable feet could easily be attached to the bottom of the drawing board.

Before mapping, we determined the species that appeared within the plot and made sure that we had enough colors of pens so that each species could be marked with a different color.

In plots that had more species than we had colored pens, we used combinations of colors and symbols.

To start mapping, first started by marking the locations of the corners of the plot on the paper map which helped us align the maps for digitization and analyses. Next, we mapped the plants starting with the most common species in the plot, mapping one species at a time. For each plant, we placed the tracing point of the pantograph arm at the location that the plant stem touches the ground (making sure not to injure or crush the surrounding vegetation). We then depressed the the pen to make a mark on the map paper. We repeated this marking for all individuals of the first species. We then added the mark and name of that species to a legend. For the next species, we replaced the color of the pen and repeated the above process for the next most populous species. We continued this process until all species were mapped onto the page.

### Map digitization protocol

To digitize each map, we started by scanning the paper map at 600 dpi (dots per inch) with a portable scanner (Doxie Go Wifi; Doxie 2016) cropping each to 4400 × 4400 pixels. Using the scanned version of each scaled map, we then digitized the maps using QGIS (QGIS Development Team, 2017; we note that a number of software packages would work for this set, including QGIS and ARCGIS). We uploaded each scanned map image as a raster file in the program. From here, we added a new ShapeFile layer on top of our raster image adding attribute data (a species code and unique individual code). We placed a ShapePoint in the ShapeFile layer were there was a dot on the scanned map (Figure 5, right panel). We did this until all points on the map had been digitized. During our point digitization, we used relative spatial distances based on a standard projection (EPSG:4326; WGS84) because we did not need specific coordinate locations for each plant. To convert distances from pixels to more meaningful measures (e.g. cm), we calculated the pixel distance of the edges and the diagonals of all plots (known values of 50 cm and 70.71 cm, respectively) and estimated an average pixel distance per centimeter scaling factor. Digitizing maps took us from 10 minutes to 5 hours to digitize depending on the density of the map (densities ranging from 50 to 800 individuals).

**Figure 5:**
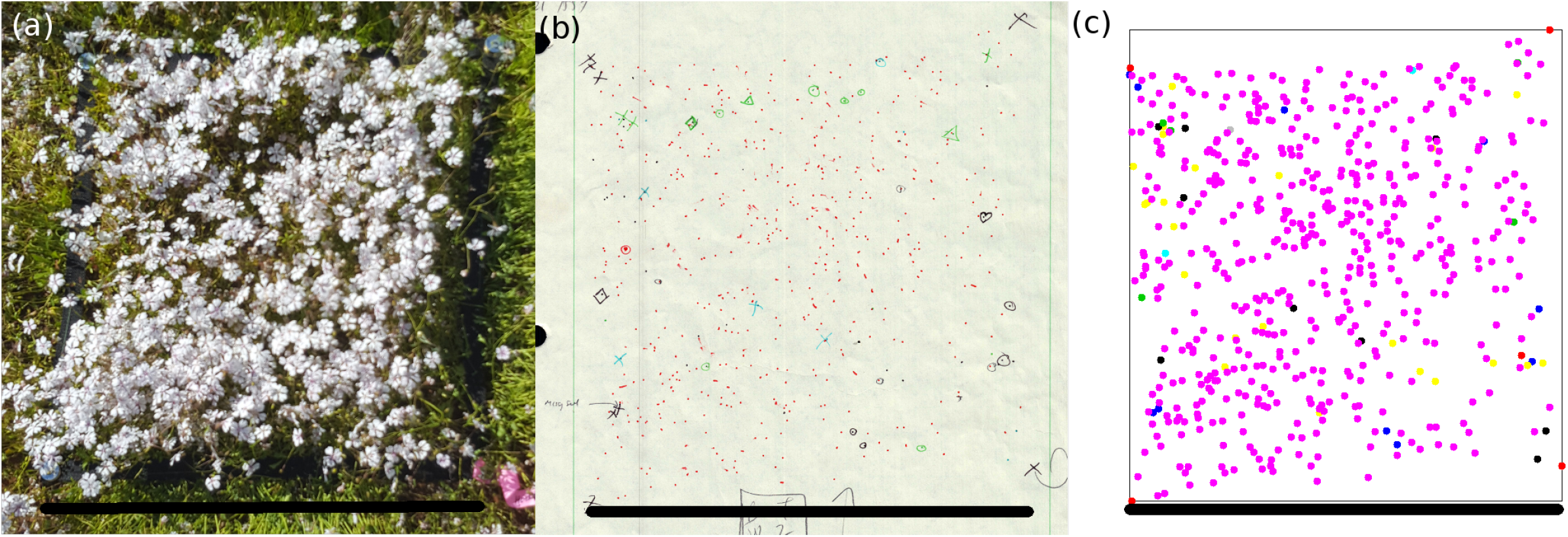
Transition from plot to data. Images of the plot in the field (a), a scanned map for the plot (b), and a digitized map of the plot (c). Black bars in each photo represent one side of the plot at 50 cm (a), 13.5 cm (b), and 4400 pixels (c).

After digitizing the maps, we loaded the data into R Statistical Software (R Core Development Team, 2017) and used various spatial packages (e.g. *spatstat* and *sp*; Baddeley & Turner 2005; Bivand *et al*. 2013) to calculate point pattern analyses, assess spatial autocorrelation in trait values, or estimate correlation weight matrices for spatial regression. The output from the digitized maps are versatile and well suited to a wide range of R-based spatial analyses, though we do not get into the details of these analyses here.

## Considerations

The pantograph arm can be made of any material (wood, plastic, or metal) depending on budget and application. Aluminum, for instance, is more expensive but is strong, light-weight, and handles moisture well. Wood is cheap and reasonably light weight but can warp easily when it gets wet and the arms are more prone to breakage. Pantographs made of wood and plastic are available for purchase, but we believe that may not be durable enough for use across multiple field seasons and various field conditions. Material and cost of the drawing board and pantograph arm should be decided based on the details of the system and project for which it is to be used. We chose our materials for sturdiness and durability to last multiple years of field work. We also considered the weight of the material because we had to ship materials to the field site and carry them up to 500 m to the plot locations within the site.

Pen type is worth considering and can depend on working conditions – waterproof, ball point, or bleed proof all have pros and cons. We used Stabilo 88 mini pens (Stablio, 2016) because they are small and thin (reducing the attachment size and weight) and they are bleed proof (making a finer point) and don’t dry out quickly if left with the cap off for a long time. Stabilo pens, however, are not waterproof and we had to be careful to keep maps dry.

The 3D pen attachment design we presented here is specific to the usage of the Stabilo Mini 88 pens (Stablio, 2016). However, a variety of pens could be used and the respective changes for the pen attachment height and diameter should be accounted for when using different pens. If you do not have access to a 3D printer, then a simple tube, such as cardboard or pvc piping just wider than the width of a pen/pencil would work. If using two people to operate the pantograph, then one person can move the arm while the other drops the pen onto the paper to make a point, this does speed up the process but is not required for the pantograph to operate properly.

There are a number of minor in-field repairs and adjustments that come up regularly with this pantograph design. For example, nylock nuts wear over time and may need to be regularly tightened or replaced so that the pantograph arm does not wobble. Users should bring extra pens, spare 3D printed pieces, springs, nylock nuts, screws, and washers into the field along with a wrench or pliers and other tools so that pieces can be quickly replaced and tightened.

Our study site was relatively flat and the PVC drawing board could be placed directly on the ground. If the intended study site has tall vegetation around the plot or uneven ground, adjustable legs or stilts can be added to the board (see Figure 2 in Hill (1920)). Another option would be to put the pantograph on a foldable camping table with legs that can be adjusted for height.

For species with a clumping forms such as perennial grasses, this pantograph can be used to trace the base or circumference of the plant. Similarly, for species with a basal rosette, both the rosette circumference and the point of ground contact can easily be traced onto the map. For species with stolons, we recommend tracing the stolon using a line or a thin polygon.

Other methods could be used and developed to digitize and process maps such as ImageJ2 (Rueden *et al*., 2017) but we decided on digitizing by hand to be able to more finely distinguish points and marks as well as add all necessary information we wanted to that point at time on digitization.

## Conclusions

In this manuscript, we re-introduced a vegetation mapping technique that can be used to identify individual plant locations in a dense herbaceous plots (Figure 5). We also present the design for a novel pen attachment to increase the efficiency of the pantograph. The pantograph has been used historically to map vegetation predominately in the Western US and the data from those plots have been applied to questions about community assembly, coexistence, and demographic research. Applications of this simple tool can also extend to spatially explicit analyses of functional traits, coexistence theory and species interactions at small spatial scales.

## Acknowledgements

We would like to thank Melissa Johnston at USDA-ARS at the CPER for her images that helped inform this paper. We would also like to thank Evan Jones and the University of Queensland Faculty of Science workshop with their help in materials acquisition and equipment fabrication.

## Appendix A: Supplemental Figures

**Figure A1:**
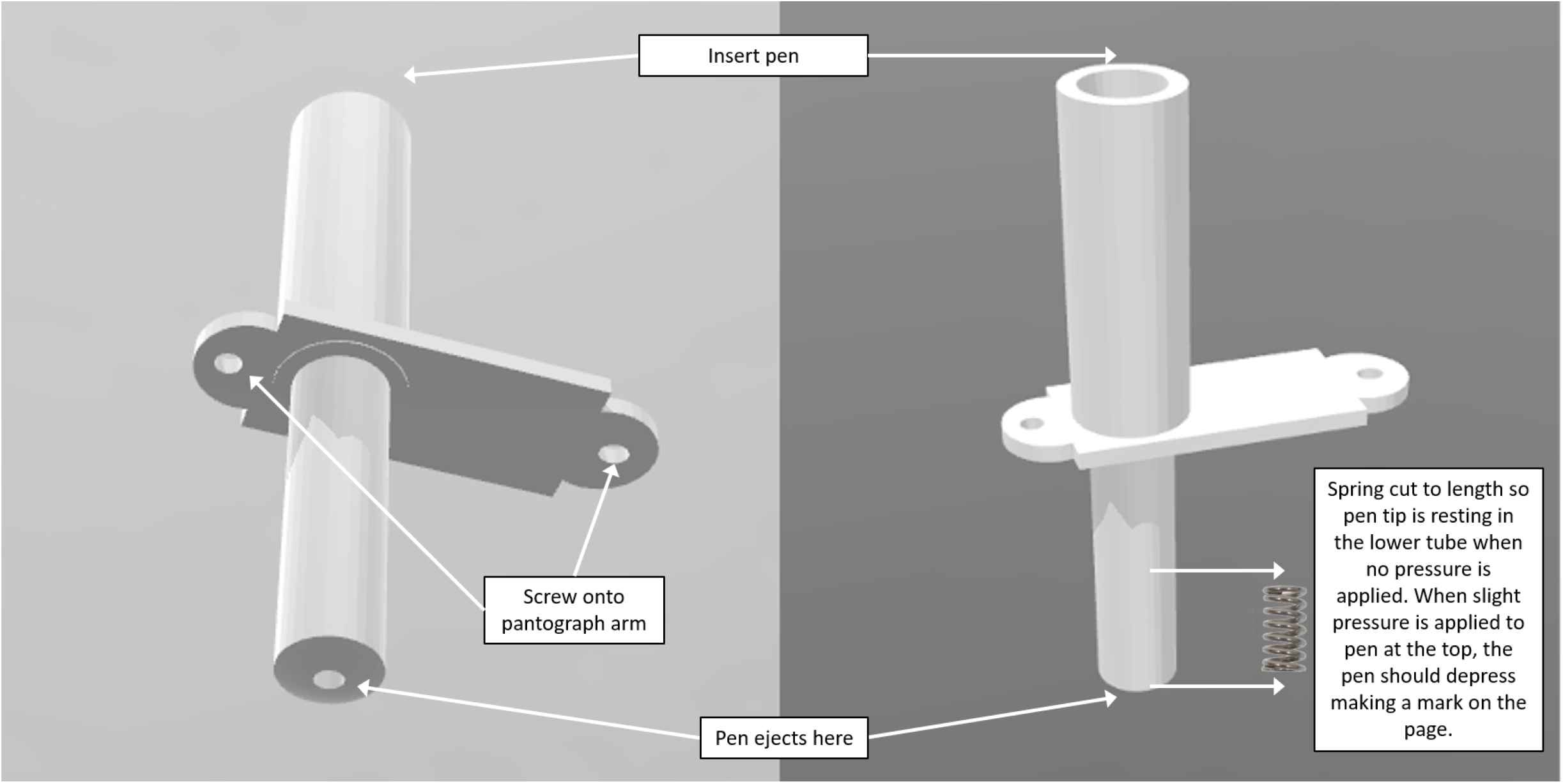
Bottom and side view of pantograph pen attachment. The left panel shows the underside of the attachment, showing where the pen would eject and mark the paper. Also shown are the attachment points. The right panel shows where to insert the pen and where in the attachment is the placement of the spring.

## Notes

### Competing Interest Statement

The authors have declared no competing interest.

